# Disarming emotional memories using Targeted Memory Reactivation during Rapid Eye Movement sleep

**DOI:** 10.1101/2024.09.25.614960

**Authors:** Viviana Greco, Tamas A. Foldes, Neil A. Harrison, Kevin Murphy, Marta Wawrzuta, Mahmoud E. A. Abdellahi, Penelope A. Lewis

**Affiliations:** Psychology Department, Cardiff University Brain Research Imaging Centre, (CUBRIC), Cardiff University, Cardiff, UK; School of Physics, Cardiff University Brain Research Imaging Centre (CUBRIC), Cardiff University, Cardiff, UK

**Keywords:** sleep, emotions, reactivation, arousal, fMRI, heart rate

## Abstract

Emotional responses are dampened across sleep, and this is thought to be mediated by neural reactivation during Rapid Eye Movement (REM) Sleep. Such reactivation can be triggered by targeted memory reactivation (TMR), a technique in which a tone previously associated with a memory during wake is re-presented during subsequent sleep. Prior work has shown that TMR in REM reduces arousal responses to negative stimuli. The present study builds on this by measuring autonomic responses and brain activity as well as behaviour. Participants rated the arousal of 48 affective images, paired with semantically matching sounds. Half of these sounds were cued during REM in the subsequent overnight sleep cycle. Participants rated the images in a Magnetic Resonance Imaging (MRI) scanner with pulse oximetry 48 hours after encoding, and again after two weeks. Results showed that TMR during REM was also associated with reduced brain activity in the two primary nodes of the Salience Network (SN): the Anterior Insula and dorsal Anterior Cingulate Cortex (dACC), as well as the orbitofrontal cortex, subgenual cingulate, and left amygdala, all of which are known to be important for emotional processing. TMR markedly reduced the emotional heart rate deceleration (HRD) response, and also reduced subjective arousal ratings for highly arousing images, while increasing ratings for less arousing images. We conclude that REM TMR can facilitate a decrease in physiological and neurological responses to arousal. These findings have potential implications for the use of TMR in treatment of depression and anxiety disorders.

**Highlights:**

- TMR in REM sleep reduces Salience Network responses to emotional pictures.
- TMR in REM sleep reduces heart rate deceleration to emotional pictures.
- TMR in REM sleep reduces subjective arousal ratings of highly arousing images.
- TMR in REM sleep provides a promising potential avenue for treatment of PTSD.

## Introduction

The unique neurological milieu of rapid eye movement (REM) sleep provides optimal conditions for the processing of emotional memories^1–7^. The “Sleep to forget, Sleep to Remember” (SFSR) hypothesis^8^ builds on this by proposing that the spontaneous reactivation of emotional memories during REM facilitates the decoupling of the affective charge from the memory.

Targeted memory reactivation (TMR) is an established technique for triggering memory reactivation during sleep. In TMR studies, stimuli, like tones or odours, that were previously associated with a newly encoded memory during wake are re-presented during sleep, prompting reactivation of the corresponding memory representation^9,10^. An extensive body of experimental work, mainly in non-REM sleep, has demonstrated the potential of this non-invasive technique to enhance the consolidation of different types of memories (see^11^ for a review). Nonetheless, the specific impacts of TMR during REM sleep on emotional reactivity remain to be explored.

We previously asked whether TMR reduces emotional reactivity by asking participants to rate emotional images for arousal both before and after the manipulation. This showed that TMR of emotionally arousing stimuli during REM, but not during slow wave sleep (SWS), led to a significant habituation of subjective arousal^12^. In the current study, we extend this REM TMR finding by examining both physiological and neural arousal responses, in addition to subjective ratings (Figure 1).

**Figure 1:**
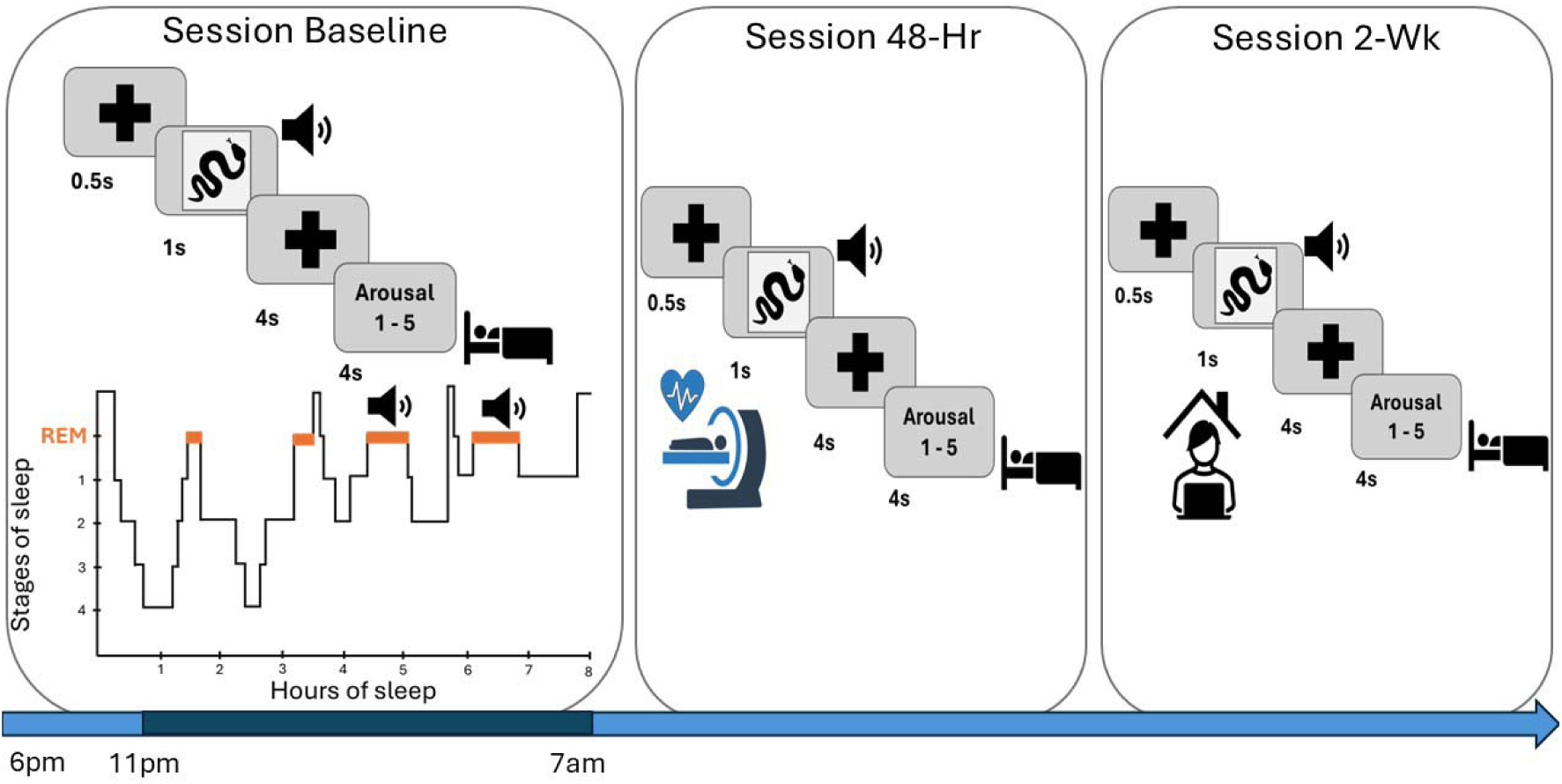
Study design. The study consisted of three sessions. At Baseline two questionnaires were first filled: the Stanford Sleepiness Scale (SSS) and the Positive and Negative Affect Schedule (PANAS). They were followed by the arousal rating task, in which participants were asked to rate 48 negative IAPS pictures -sound pairs on a 5-point rating scale of arousal (1 = less arousing, 5 = more arousing). After filling out the PANAS a second time, participants were wired-up for EEG. During the night, tones associated with half of the arousing stimuli were played in random order during REM sleep. The following morning, participants filled a sleep quality questionnaire. At the second session (Session 48-H) and the third session (Session 2-Wk) had the same structure: after filling the SSS and the PANAS, participants performed the arousal rating task. The PANAS was administered again as soon as they finished the task. Session 48-Hr was performed in the MRI scanner while heart rate deceleration (HRD) was recorded; it occurred 48 h after S1. Session 2-Wk was performed online (2 weeks after Baseline).

We used functional magnetic resonance imaging (fMRI) to examine brain activity, focusing on regions known to be involved in the processing and regulation of emotions, namely the amygdala, insula, orbitofrontal cortex (OFC), subgenual anterior cingulate cortex (sgACC). These regions have been extensively studied and are recognized as key components of the neural circuitry underlying emotional experiences. Previous studies have reported alterations in activations here as a result of sleep^6,13^ or emotional reactivity^14–17^. Furthermore, dysfunctions within these regions have been associated with psychiatric disorders such as depression or post-traumatic stress disorder(PTSD)^18–21^.

We used heart rate deceleration (HRD) as a physiological marker of autonomic arousal. HRD, which reflects the parasympathetic orienting response, has been shown to map onto the affective tone of a stimulus, with greater deceleration indicating higher levels of arousal^22,23^. Prior work has discussed different effects of sleep on the parasympathetic aspects of emotional arousal, with studies showing either a decrease^24,25^ or a perseveration of the HRD in response to emotional stimuli^26–28^. No study has specifically explored the response of HRD to TMR during REM.

Drawing on our own previous findings^12^, and supported by evidence indicating that REM sleep can provide optimal conditions for the processing of emotional memories^7,8,29^ and by studies suggesting that REM TMR may impact upon arousal responses^6,30,31^, we predicted that our manipulation would result in diminished activity in the brain’s arousal system, decreased heart rate responses, and reduction in subjective arousal during the processing of emotional images.

## Results and Discussion

### Sleep characteristics and EEG analysis

Participants obtained an average of 528.91 min total sleep time (TST) (+/- 37.38), with an average of 95.09 min of REM (+/- 27.42 min, see Table S1 for full details of sleep). EEG analysis confirmed that TMR cues were processed by the brain since TMR onset is followed by an increase in beta band (12.5-30 Hz) compared to baseline (see figure S1B). This starts around one second after TMR (corrected with cluster-based permutation, n = 16, p = 0.0079). ERP analysis showed an amplitude increase just before 500ms after the TMR onset, followed by a decrease 500ms after the cue (S1B). These findings are in keeping with other TMR studies which used baseline correction rather than control tones in REM^32^ and NREM^6,33^, both of which show similar pattern in the ERP and time-frequency response. Notably, however, the absence of a control tones (not associated with a memory) makes it difficult to say whether the elicited responses are due to memory reactivation or are instead related only to the sound.

### fMRI

To determine whether REM TMR led to a decrease in neural arousal responses in the brain, we compared fMRI responses to Uncued > Cued picture-sound pairs. We first tested this using a whole-brain corrected analysis of grey matter which revealed two strong decreases of response in dorsal anterior cingulate cortex (dACC) (349 voxels and 159 voxels), with no other clusters surviving (Figure 2a, Table 1). Next, we tested the same comparison in our insula ROI and found reduced responses in the anterior portion of insula (Figure 2b, Table 1), FWE corrected at p<0.05.

**Figure 2.**
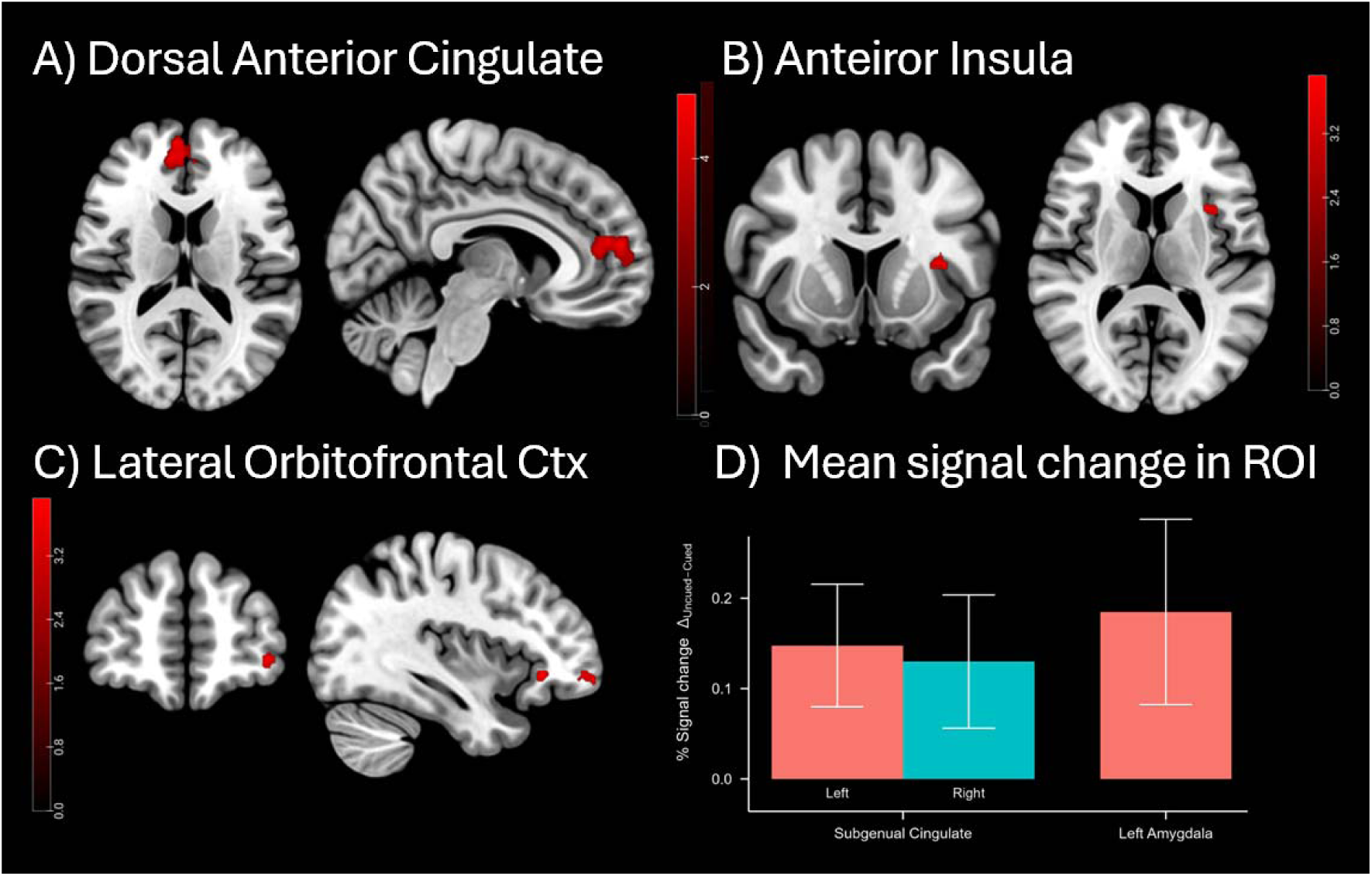
Functional activity in response to Uncued > Cued contrast on negative picture-sound pairs in Session 48-Hr. A-C) Cluster corrected responses at pFWE < 0.05, cluster-level corrected. Results are overlaid on a skull stripped MNI ICBM152 T1 template. OFC = orbitofrontal cortex. D) Percent signal change (Uncued > Cued) in sgACC (R and L) and L Amygdala. Bars represent mean percent signal change in the Uncued minus Cued contrast, with error bars indicating standard error of the mean. Positive values indicate less activation in the Cued condition compared to the Uncued condition.

**Table 1.** Functional results. Peak Z-values and corresponding MNI coordinates for regions showing activation in the contrast Uncued > Cued with the inclusion threshold of one-tailed p<0.001 and cluster correction of p<0.05.

| Brain region | No. voxels | Peak Z-value | MNI x, y, z (mm) |
| --- | --- | --- | --- |
| Insula | 28 | 4.01 | -33, 14, 14 |
| Orbitofrontal Cortex | 39 | 4.29 | -37, 60, -9 |
|  | 17 | 3.93 | -37, 32, -7 |
| Dosal Anterior | 349 | 5.13 | 5, 56, 14 |
| Cingulate | 121 | 4.5 | -39, 44, 2 |

Anterior insula and dACC are the two primary nodes of the brain’s Salience Network (SN)^30,31^, so a reduction in responsivity in these regions after REM TMR suggests that our manipulation could lead to downscaling of a more generalised salience response. The SN helps the brain to identify important stimuli, and to coordinate resources in response to these stimuli, for instance by switching between the Central Executive Network and the Default Mode Network. Within this network, the frontal insula is an afferent hub for detecting autonomic feedback, while the dACC is the efferent hub, important for generating responses. Together, these regions are thought to process salient stimuli, determine their importance, and select responses^30,34^, for a review see^31^.

The SN influences physiological arousal via connections to the amygdala which recognizes threats and recruits brain structures to respond to threat^17,31^. Examination of our amygdala ROI showed a reduction in response in just one voxel of left amygdala at p < 0.001 uncorrected, (x = -26, y = 1, z = 16), however this did not survive FWE correction. To check for a more diffuse response across this structure, we next examined mean percent signal change in our L amygdala ROI and found a significant effect (M = 0.185%, SD = 0.436%), t(17) = 1.80, p = .045, see Figure 2d. This suggests that our TMR manipulation leads to a dispersed change across the left amygdala, rather than being apparent in a strong cluster. Such a change in reactivity is in keeping with the literature since amygdala responses to arousal have already been shown to be modulated by REM TMR^6^ although the effect is disrupted when REM is extremely disturbed. These findings might indicate that REM TMR can reduce the extent to which a negative stimulus is perceived as a threat.

Next, we examined our ROI in orbitofrontal cortex, finding a strong reduction in right lateral OFC at p<0.05 FEW corrected, (figure 2c). OFC plays a critical role in representing the reward value associated with a range of stimuli and outcomes. It encodes the emotional and affective significance of different inputs, thus contributing to the modulation of emotional responses^14^. The significance of OFC in shaping emotional experiences and behavioural responses becomes even more apparent when we consider outputs to regions such as the dACC and insula since these allow reward value representations generated by the OFC to feed into the SN and contribute to the complexity of our emotional experiences and associated behaviours^14,17^.

Our final ROI was the subgenual cingulate, a region which is associated with psychiatric disorders such as depression or PTSD^18,19^. Here, as in amygdala, we observed uncorrected responses (just one voxel in each case). Because these did not survive FWE correction, we examined mean signal change for each area and found significant results in both hemispheres: (left sgACC: M = 0.148%, SD = 0.288%), t(17) = 2.18, p = .022, right sgACC: (M = 0.130%, SD = 0.313%), t(17) = 1.76, p = .048, figure 3D.

**Figure 3.**
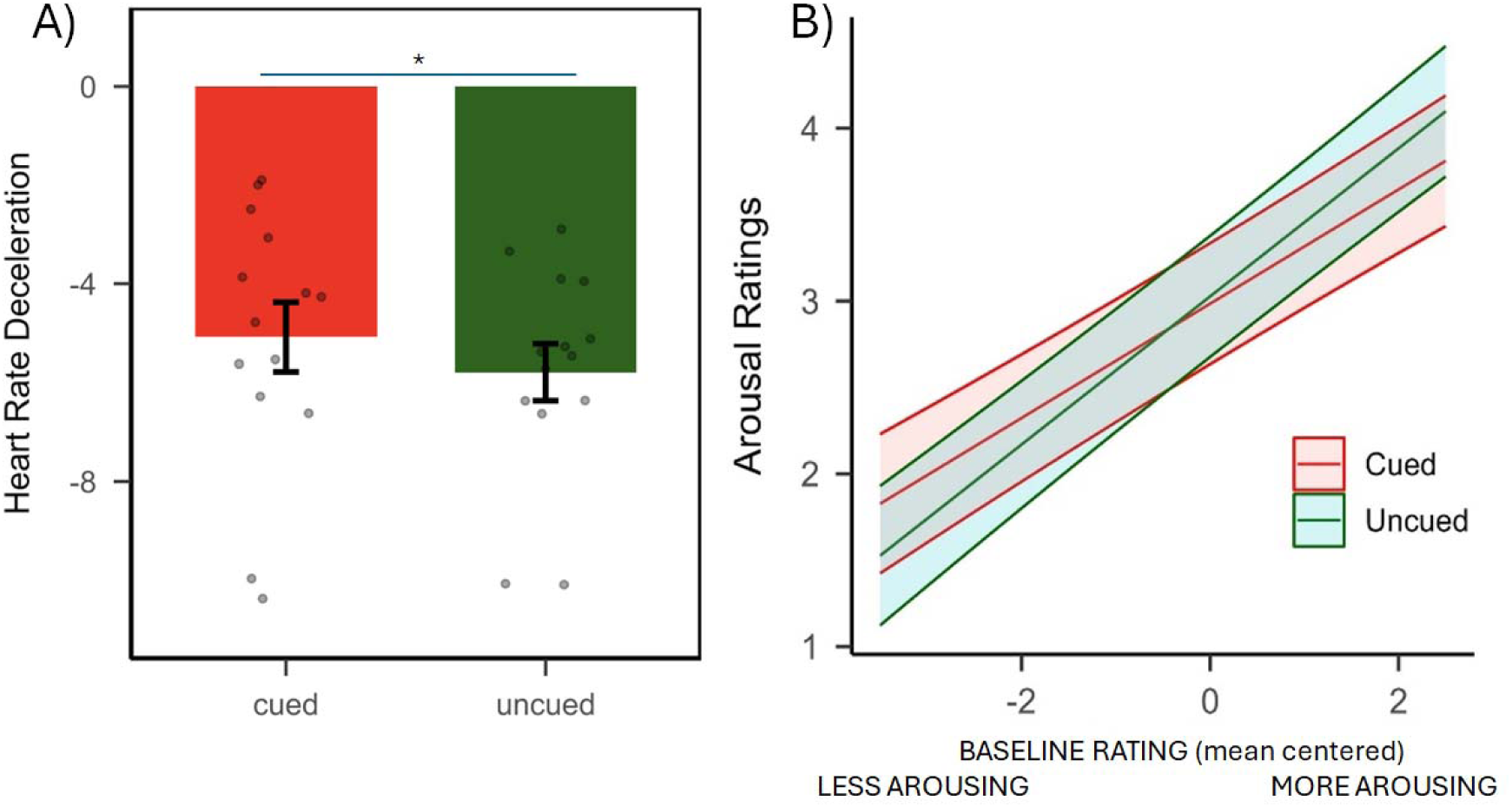
**(a) Post-manipulation Behavioural data (Sessions 48-H and 2-Wk):** Model predicted interaction between Cueing and mean-centered Baseline. Shadowed areas represent 95% Confidence Intervals. **(b) Heart rate deceleration**: HR responses for the negative Cued (red) and Uncued (green) image-sound pairs at Session 48-H. *P = <0.05. Data are shown as means (±SEM). Grey dots represent individual participants.

### Heart rate deceleration (HRD)

To index the impact of the TMR manipulation on autonomic responses to negative images, we compared heart rate deceleration between Cued and Uncued images in the second Session 48-H. This revealed greater deceleration for Uncued stimuli (M ± SE Cued= -4.53 ± 0.68; M ± SE Uncued = -5.24 ± 0.98; paired t-test: t_13_ = 2.51; p = 0.02), indicating a stronger emotional reactivity to these Uncued images (Figure 3a). No significant correlations were found between HRD, sleep, behavioural results, or functional parameter estimates (all p_adj_ > 0.05, see Supplemental Material Table S2 and S3).

Our observation of depotentiated visceral reactivity to stimuli that were Cued overnight is in good keeping with the observed downregulation of the Salience Network. The majority of studies indicating a sleep-dependent preservation of physiological arousal in response to negative stimuli have investigated the effect of either a nap or a single night of sleep^26–28^. One study of how sleep modulates affective reactivity observed that HRD was preserved in the short term, but reduced after a week, suggesting that time might play an important role in the modulation of emotional strength^24^. Because we observed a reduction in HRD 48h after the first exposure to the task followed by the TMR manipulation we speculate that the TMR may have speeded up this time dependent decrease.

### Subjective Arousal Ratings

To investigate the effects of REM TMR on subjective arousal ratings we used a linear mixed model (LMM) with Cueing (Cued and Uncued), group mean centred ratings at Baseline, and Session (48-H and 2-Wk) as fixed effects (formula: rating ∼ Cueing * Session + Cueing * Baseline). Both participants and items were included as random effects (formula: ∼ 1 | Participants, ∼ 1 | Items). There was no effect of cueing (M = −0.04, 95% CI [−0.15, 0.07], p = 0.45, but the interaction between Cueing and Baseline was significant (M = −0.10, 95% CI [−0.17, −0.02], p = 0.010) (Figure 3b), see Table S2a for full results). Thus, when baseline arousal ratings were taken into account, REM TMR led to decreases in arousal ratings of pictures that had been rated as very arousing at baseline while also leading to an increase in arousal ratings for pictures rated as less arousing at baseline. This pattern is consistent with the work of Pereira et al. (2022) who observed that Slow Wave Sleep (SWS) TMR of emotional material reduced responses in the orbitofrontal cortex for negative items, while simultaneously increasing them for neutral items^35^. We speculate that this could be due to a generalization effect of TMR, where the manipulation leads to a reduced emotional intensity of stimuli characterized by higher initial arousing levels, while simultaneously increasing the intensity of stimuli characterized by lower initial arousal levels.

Notably, Session also had a significant negative effect (M = −0.29, 95% CI [−0.41, −0.17], p < 0.001), suggesting that ratings decreased across sessions, but the interaction between Cueing and Session was not significant (M = 0.05, 95% CI [−0.12, 0.21], p = 0.564), suggesting that the effect of cueing did not vary significantly across sessions. Post-hoc t-tests (Table S2B) revealed that that cueing tended to decrease arousal ratings for items that were rated as higher than average arousal at baseline, while simultaneously increasing ratings for items rated lower than average arousal at baseline (Figure 3a). Descriptive statistics are reported in Table S3.

### Synthesis

Our observation that REM TMR leads to reduced arousal related responses in the salience network as well as the amygdala, orbitofrontal cortex, and subgenual cingulate combines with our behavioural and autonomic results to suggest that REM reactivation can somehow reduce the extent to which an emotional stimulus elicits arousal, and this is true not only subjectively, but also in terms of autonomic and neural responsivity. Our current findings join prior research showing promising evidence for a role of memory reactivation during REM sleep in decreasing the affective tone associated with negative experiences. For instance, Wassing and colleagues induced the self-conscious emotion of shame in volunteers suffering from insomnia to explore the impact of disrupted REM sleep on emotional distress^36^. Their findings indicate that discontinuities in REM can prevent the brain from processing and reducing emotional distress as reflected by continuous amygdala reactivity. Another recent study showed that TMR during REM of imagery rehearsal therapy for two consecutive weeks reduced the frequency of nightmares while promoting more positive dream emotions ^37^.

Overall, our findings support the possibility that targeted reactivation of emotionally arousing memories in REM could potentially offer a way to make these memories less upsetting. As such, our method could lead to clinically important opportunities for the early treatment of psychiatric disorders such as depression and post-traumatic stress disorder PTSD.

## Acknowledgements

We thank Sofia Periera, Sophie Smith, and Caterina Leitner for collaboration at the early and late phases of the study. This work was funded by the ERC consolidator grant SolutionSleep to Lewis P.A. Murphey K was funded by The Wellcome Trust [WT224267]. For the purpose of open access, the author has applied a CC BY public copyright licence to any Author Accepted Manuscript version arising from this submission.

## Author contributions

Greco V., Pereira I.R.S., Harrison N. and Lewis A.P. designed the study. Greco V. and Pereira I.R.S. performed the experiment. Greco V. and Foldes T. analysed data. Harrison NA and Murphy KA advised on aspects of data analysis. Foldes T. wrote the fMRI pipeline and contributed to interpreting the results. Wawrzuta M and Abdellahi M. analysed the EEG data. Lewis P. A. and Greco V. wrote the manuscript. Foldes T. and Lewis P.A. contributed to editing the manuscript.

## Star Methods

### Participants

Twenty-three right-handed, non-smoking healthy volunteers (14 females, age range: 20 – 33 years, mean ± SD: 23.61 ± 3.92) were recruited for this study, which was approved by the Ethics Committee of the School of Psychology at Cardiff University. A pre-screening questionnaire was used to ensure that participants were fluent in English, had normal or corrected to normal vision, no previous history of physical, psychological, neurological, or sleep disorders and no hearing impairments. Participants were required to be right-handed and to not regularly take any psychologically active medication or substance directly or indirectly affecting sleep quality. They agreed to abstain from alcohol 24 hours prior to each experimental session and from caffeine and other psychologically active food from 12 hours prior. Participants were also asked to refrain from engaging in intense physical activities during the period of the study. Further criteria of exclusion included a habit of daytime napping, a non-regular sleep-wake rhythm, engaging in nightshift work, cross-continental travel in the two months before the study or having such plans during the experimental weeks. Additionally, to ensure that participants did not experience negative emotional stress over the week before starting the experiment, they were asked to complete the Depression, Anxiety and Stress Scale as inclusion criteria (DASS-42, normal scores: Depression (D) ≤ 9; Anxiety (A) ≤ 7; Stress (S) ≤ 14)) ^38^. All participants gave written informed consent and received monetary compensation for their participation. Five participants were excluded from all analyses due to: voluntary withdrawal (n = 4) or technical issues (n = 1) and three participants were unable to complete the online follow-up. Hence, the final dataset included 18 participants (11 females, age range: 20 – 30 years, mean ± SD: 23.61 ± 3.56) at Baseline and 48-h, and n = 15 in 10-days (S3).

### Arousal rating task

Participants viewed 48 standardized negative images selected from the International Affective Picture System ^39^ (see Supplemental Material Table S5). Each image was converted to greyscale and matched in luminance and resolution (height = 600 px; width = 800 px) using the SHINE toolbox ^40^ in MATLAB 2007a. Each image was rated using a 5-point arousal scale, corresponding to increased emotional intensity (i.e. 1 = less arousing, 5 = more arousing), and paired with a semantically related sound obtained from the International Affective Digitized Sounds database (IADS; ^39^). Participants were instructed to rate each picture-sound pair along arousal dimension.

Each trial consisted of a fixation cross (500ms), picture and sound presentation (1 s), a blank screen (4 s), the arousal rating (4 s) and the inter-trial interval (jittered: 3.5 – 4.5 – 5.5 – 6.5 s). Sounds were 400ms long. In order to match the duration of the picture presentation on the screen (1s) with the duration of the sounds, the 400ms sounds were repeated twice with a 200ms gap in between the two presentations: 400ms – 200ms gap – 400ms. Audacity software (www.audacityteam.org) was used to modify the length of sounds.

In Session 1 only, the arousal rating task was preceded by a practice round and followed by a forced-choice task. The practice round aimed to let participants familiarize with the rating scale. It consisted of four neutral IAPS pictures paired with semantically related neutral sounds taken from the IADS. The forced-choice task aimed to assess whether participants had learned the associations between images and sounds. For each trial participants had to choose which of the four IAPS images displayed was semantically related with the sound. This task was repeated until participants reached 75% accuracy. Feedback with the correct answer was presented for 1.5 s.

### Study design and procedure

The study consisted of three sessions (Figure 1), all scheduled for the same time in the evening (∼6pm). For all sessions, before and after performing the arousal rating task participants completed the Positive and Negative Affect Schedule (PANAS) scale ^41^ to evaluate their mood. The Stanford Sleepiness Scale ^42^ was administered at the beginning of each experimental session to determine participants’ level of alertness.

S1 lasted approximately 2h and progressed as follows: participants completed the arousal rating task, then changed into their sleepwear, were fitted for polysomnography (PSG) recording and went to bed at around 11:30 pm while brown noise was delivered throughout the night to minimize noise-induced arousals. For the TMR protocol acoustic stimuli semantically related to the IAPS pictures were replayed during REM sleep to trigger reactivation of negative emotions. Participants were woken up after 7-8 hours of sleep. After removing the electrodes and before leaving the lab, they were asked to rate their sleep quality and whether they heard any sounds during the night with an adapted and translated version of a German sleep quality questionnaire^43^. Participants were asked to come back to the lab 48-hours later for Session 48-H during which the arousal rating task was performed in a 3T Siemens MRI scanner during fMRI acquisition while heart rate deceleration (HRD) was recorded. Session 2-Wk (2 weeks after Baseline), the follow-up session, was performed online and lasted ∼40 minutes.

both the lab and the MR scanner, the task was presented using PsychoPy3 Experiment Runner (v2020.1.3)^44^. The SSS and the PANAS questionnaires were executed using MATLAB (The MathWorks Inc., Natick, MA, 2000) and Psychophysics Toolbox Version 3^45^, except for the sleep quality questionnaire, completed with pen and paper. In S3 the behavioural task was administered through the Pavlovia online platform (https://pavlovia.org/) and the SSS and PANAS questionnaire were distributed via Qualtrics software (Qualtrics, Provo, UT, USA. https://www.qualtrics.com).

### Questionnaires

The SSS is used to provide a subjective indication of sleepiness, with participants rating their current state on a 7-point Likert scale, where 1 is most alert and 7 is least alert^42^.

The PANAS scale^41^ is a self-report measure composed of two subscales designed to assess individuals’ levels of positive and negative affect. Each subscale is composed of 10 Likert-type format items ranging from 1 (vey slightly/not at all) to 5 (very much).

### PSG data acquisition

Standard polysomnography consisting of electroencephalography (EEG), left and right electromyography (EMG) electrodes placed on the chin, left and right electrooculography (EOG) electrodes placed below and above the eyes, was continuously recorded using passive Ag/AgCl electrodes and collected with a BrainAmp DC Amplifier (Brain Products GmbH, Gilching, Germany). According to the international 10–20 system, six EEG electrodes were positioned on the scalp (F3, F4, C3, C4, O1 and O2) and we further attached one ground electrode to the forehead. All electrodes were referenced to the mean of the left and right mastoid electrodes applied behind the left and right ear. Impedances were maintained below 5 kΩ for each scalp electrode, and below 10 kΩ for each face electrode. Electrodes were applied with Ten20 conductive paste (Weaver & Co., Aurora, USA) on sites cleaned with NuPrep exfoliating gel (Weaver & Co., Aurora, USA). Data were recorded using BrainVision Recorder software (Brain Products GmbH), sampled at 500 Hz and saved without further filtering.

### TMR during REM sleep

Acoustic stimuli which had been paired with pictures during wake were replayed to the participants during stable REM sleep, as assessed with standard AASM criteria ^46^. The TMR protocol was executed using MATLAB 2016b and Cogent 2000 and it consisted of the presentation of 24 Cued sounds (400ms duration) repeatedly presented 20 times each (20 loops), with an inter-trial interval jittered between 2, 2.5, 3, 3.5 and 4 seconds. Volume was adjusted for each participant to make sure that the sounds did not wake them up and to prevent arousals. Cueing was paused immediately when any sign of arousal was showed or when participants left the relevant sleep stage and resumed only when stable REM sleep was observed. Notably, post-hoc scoring verified that all TMR cues were delivered in REM for 17 of the participants for whom data were analysed. In one participant 24 out of the 480 cues were erroneously delivered in NREM.

### MRI data acquisition

Magnetic resonance imaging (MRI) data were obtained at Cardiff University Brain Imaging Centre (CUBRIC), using a Siemens Magnetom Prisma 3T scanner with a 32-channel head coil. Functional images were acquired with a T2*-weighted echo-planar imaging (EPI) sequence (repetition time (TR) = 2000 ms; echo time (TE) = 30 ms; FA = 75°; bandwidth 2442 Hz/Pixel, field of view (FoV) = 224 mm^2^; voxel-size = 3.5 mm^3^; slice thickness = 3.5 mm; 37 slices with a ∼25° axial-to-coronal tilt from the anterior – posterior commissure (AC-PC) line and interleaved slice acquisition; parallel acquisition technique (PAT) with in-plane acceleration factor 2 (GRAPPA), anterior-to-posterior phase-encoding direction). To correct for distortions in the fMRI data caused by magnetic field inhomogeneities, B0-fieldmap was acquired (TR = 1000 ms; TE1 = 4.92 ms; TE2 = 7.38 ms; FA = 75°; bandwidth 290 Hz/Pixel; FoV = 224 mm^2^; voxel-size = 3.5 mm^3^; slice thickness = 3.5 mm; interleaved slice acquisition; anterior-to-posterior phase-encoding direction).T1-weighted structural images were obtained using a 3D magnetization-prepared rapid-acquisition gradient echoes (MPRAGE) sequence (TR = 2300 ms; TE = 3.06 ms; FA = 9°; bandwidth 230 Hz/Pixel, FoV = 256 mm^2^, voxel-size = 1 mm^3^, slice thickness = 1 mm, parallel acquisition technique (PAT) with in-plane acceleration factor 2 (GRAPPA), anterior-to-posterior phase-encoding direction).

## Data analysis

### Behavioural data analysis

Differences on arousal ratings between Cued and Uncued items were assessed using a linear mixed effects models implemented in the lme4 package^47^.

To identify the contribution of Cueing on arousal ratings across time, we first fitted a model that included Cueing (two levels: Cued and Uncued), Session (two levels: S2 and S3) and their interaction as fixed effects, and participants and items as random effects.

**Model 1 formula**: Rating ∼ Cueing * Session + 1 | Participants, ∼ 1 | Items.

Next, we introduced the baseline variable into the model, representing the ratings provided by participants in Session 1, before any experimental manipulation.

**Model 2 formula**: Rating ∼ Cueing * Session + Baseline + 1 | Participants, ∼ 1 | Items.

We employed a *group mean centering* (GMC) for the baseline values by subtracting each individual baseline rating from their overall score^48^. By adopting this approach, we ensured a more accurate evaluation of the changes occurring within participant and mitigated the influence of divergent initial rating levels between individuals. Moreover, we addressed the potential multicollinearity among predictor variables in the model^49^.

Finally, we added an interaction term between cueing and group mean centered baseline to examine whether the effect of cueing on arousal ratings varied depending on the participants baseline level.

**Model 3 formula**: rating ∼ Cueing * Session + Cueing * Baseline + 1 | participants, ∼ 1 | items.

To determine statistical significance, we conducted a likelihood ratio test (LRT) in which we compared all the three models. The LRT yielded a substantial improvement in the model fit (*χ*2(1) = 6.60, p = 0.01), thus we will report our analysis based on this third model. For a model comparison analysis see Supplemental Material Table S4.

We used R (Rstudio Team (2022), www.R-project.org) and the R-packages *lme4* and *emmeans* for all our statistical analyses^47,50^. Figures were created using ggplot2 package^51^.

Finally, were interested in determining whether time spent in REM or SWS modulated the effects of TMR. Unfortunately, the sleep data were missing for two participants due to technical problems, and thus including sleep data in the analysis reduced our sample size. We nevertheless examined the subset of data with complete sleep information to determine whether adding these covariates would have improved model fit. For SWS, a likelihood ratio test comparing the original model to a model including SWS time as a covariate revealed no significant improvement in model fit, χ²(1) = 0.32, p = .574. Similarly, adding REM sleep time as a covariate did not significantly improve model fit compared to the original model, χ²(1) = 0.09, p = .759. These results suggest that the inclusion of SWS and REM sleep time as covariates would not have substantially altered the findings reported in the main analysis.

### EEG data analysis

PSG recordings were manually scored in 30s epochs by two trained independent sleep scorers, according to the standard AASM manual^46^. Each EEG recording was scored using a publicly available interface (https://github.com/mnavarretem/psgScore). From the scored sleep stages, the following sleep macrostructure parameters were calculated: (1) total sleep time (TST, min) as the total time in any sleep stages other than wake; (2) time spent in each sleep stage; (3) percentage of time spent in each sleep stage, calculated as the time in the respective sleep stage over TST. Data from N = 2 participants was excluded due to recording issues. Sleep parameters are reported in Table 1.

### EEG cleaning

EEG cleaning consisted of filtering and rejection of outliers based on statistical measures. EEG cleaning began with band-pass filtering (0.1 to 30Hz) and band-stop filtering (50Hz). EEGs were segmented into 3-second trials (0.5 sec. pre-stimulus and 2.5sec. post-stimulus). We removed trials representing outliers based on statistical measures (variance, max, min) extracted for every trial and every channel. A trial was considered as an outlier if its statistical measure exceeded the third quartile + (the interquartile range *1.5) or was below the first quartile - (the interquartile range*1.5) in more than 25% of channels. This was done for all mentioned statistical measures. If a trial was marked as an outlier for less than 25% of channels it was interpolated using neighbouring channels with triangulation method in Fieldtrip, otherwise, it was removed. Trials were then visually inspected, and any remaining artifacts were removed.

### Time-frequency representation and ERP analysis

We performed time-frequency decomposition in a similar way to that used in prior reports^52,53^. We used a hanning taper with 5 cycles that was convolved with the signals. We used 0.5 Hz frequency steps and 5 ms time steps. Power values are shown in the range of 7–30 Hz, Figure s1a. We also used a baseline of –400 ms to 0 ms relative to the onset of the TMR. The reported values represent the percentage of power change from baseline. Missing values at the edges are caused by using 5 cycles of the estimated frequency to have an adaptive window as a function of frequency. The shown plots are the grand average from all participants and all channels. For the ERP analysis, we identified a baseline period of –400 ms to 0 ms and again we report the grand average from all participants and all channels. Small values of amplitudes shown in the ERP plot (Figure 4a) are caused by the smoothing that happened as a result of averaging many trials, participants and channels, thus small shifts between values will make amplitude values smaller as shown.

### Correction for multiple comparisons

Time frequency decomposition was compared to baseline and was corrected for multiple comparisons using cluster-based permutation in Fieldtrip^54^ and lively vectors (lv)^55^ which have the same results. For cluster-based permutation, Monte Carlo was used with a sample-specific test statistic threshold = 0.05, permutation test threshold for clusters = 0.05, and 10,000 permutations. The correction window was the whole length of the plot after removing missing values. Plots of ERP analysis and time-frequency analysis and cluster-based permutation were built with lively vectors (lv)^55^.

### MRI data analysis

Image data preparation, preprocessing, and statistical analysis were performed using fMRIPrep 20.2.7 (RRID:SCR_016216 ^56^) which is based on Nipype 1.7.0 (RRID:SCR_002502 ^57^). Functional data were preprocessed in the following way: (1) a B0-nonuniformity map correction (or field map); (2) co-registration to the participants’ T1-weighted anatomical scan using rigid-body model; (3) motion correction (transformation matrices, and six corresponding rotation and translation parameters); (4) slice-time correction to 0.481; (5) spatial normalization to Montreal Neurological Institute brain (MNI space); (6) resampling to a voxel size of 2x2x2 mm using cubic interpolation; (7) smoothing using a Gaussian kernel with a full-width half maximum (FWHM) of 6 x 6 x 6 mm.

### First and second level analysis

Subject-level analysis was performed using a general linear model constructed separately for each participant. The design matrix included two regressors: Cued and Uncued picture-sound pairs. Each regressor was convolved with a canonical haemodynamic response function using the default Glover HRF in Nilearn. Additionally, six affine motion correction regressors estimated during realignment (translations in x, y, z directions and rotations around x, y, and z axes) were included as non-convolved regressors of no interest in the matrix.

To mitigate the effects of excessive motion during the fMRI scan, we employed scrubbing as a denoising approach^58–60^. Scrubbing involved identifying volumes in the fMRI data that exhibit high motion and excluding them from statistical analysis. Frame displacement (FD), a measure of head motion between consecutive frames in fMRI, was used to define excessive motion^60^. Volumes exceeding a specified threshold (0.5, as suggested by^60^), were considered to have excessive motion and were excluded or “scrubbed” from further analysis.

The effect of cueing in REM sleep was estimated using a one-tailed t-test for Uncued>Cued. Individual contrast images resulting from the first-level analysis were carried forward to the second-level one-way t-tests.

A-priori defined ROIs consisted of the insula, sgACC, OFC and amygdala. These regions were selected based on previous findings that reported activations (or de-activations) in these regions due to sleep or emotional^6,7,14,61^ and their involvement in psychiatric disorders^19,62^. ROIs were created using the integrated Automated Anatomical Labeling (AAL) atlas^63^ in the Wake Forest University Pick Atlas toolbox (http://fmri.wfubmc.edu/software/PickAtlas) and the automated anatomical labelling atlas 3 template (AAL3^14^) was used to define the sgACC. The masks were thresholded at 0.1. In addition, we included a whole brain gray matter (GM) mask thresholded at 0.1.

To control for multiple comparisons, we performed cluster-level corrections. This was accomplished using the 3dttest++ function in the Analysis of Functional NeuroImages (AFNI) software suite^64,65^, employing the ClustSim option for Monte Carlo simulations. A cluster-defining threshold (CDT) of p < 0.001 was set to identify potential clusters showing a significant effect. The ClustSim option generated a distribution of cluster sizes under the null hypothesis, allowing us to determine a cluster-size threshold corresponding to a family-wise error (FWE) corrected p-value of less than 0.05.

An additional region of interest (ROI) analysis was performed to investigate the mean activation within the left and right subgenual cingulate cortex (sgACC) and the left amygdala. For each participant, contrast images representing the difference in activation between the Uncued and Cued conditions (Uncued-Cued) were generated from the first-level fMRI analysis. These contrast images were then masked with the sgACC and left amygdala ROIs using a custom Python script leveraging the NiBabel and NumPy libraries. The script extracted all voxel values within each ROI for each participant and calculated the mean activation within each mask. This resulted in a single mean activation value for each participant and ROI, representing the average difference in activation between the Uncued and Cued conditions within that region. To examine whether cueing led to deactivation within these ROIs, a one-tailed one-sample t-test was performed on the mean activation values at the group level, testing the hypothesis that the mean activation was significantly higher than zero.

### Heart rate data analysis

At Session 48-H (when the task was performed in the MR scanner), heart rate was acquired with a pulse oximetry sensor provided with the Siemens Physiological Monitoring Unit and attached to the ring finger of the non-dominant hand. R components of the QRS complexes were marked using custom made script in Matlab 2019a and subsequently interpolated at 1000 Hz. HRD was computed as the maximum R-R interval deceleration in the 5 s interval following each picture onset, subtracted from the mean R-R interval during the 1.5 s baseline period before each picture onset. Due to technical difficulties (high presence of motion artifacts n = 3 and poor sensor placement n = 1), only data from N = 14 participants were analysed.

To compare HRD between Cued and Uncued stimuli we used paired-samples t-test (Gaussian distribution). Correlations between HRD, behavioural measures, parameter estimates for our ROIs in each subject and EEG results were assessed with Pearson’s correlation or Spearman’s Rho (depending on the Shapiro-Wilk test result) using cor.test() function in the R environment. False discovery rate (FDR) correction was used to correct for multiple correlations (q < 0.05) ^66^.

