## Supplementary tables and figures for "Disarming emotional memories using Targeted Memory Reactivation during Rapid Eye Movement sleep"

*Lewis A. Penelope

### Supplemental figures

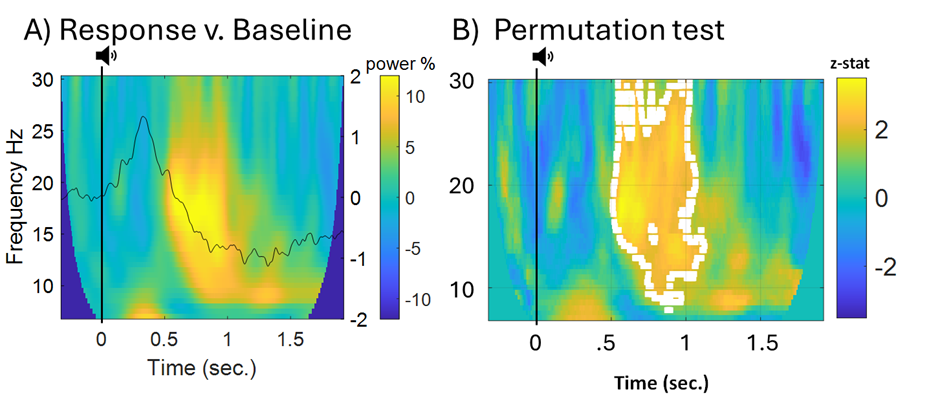

**Figure S1. ERP and time-frequency analyses**, A) ERP analysis using baseline duration of 400 ms pre-stimulus and time-frequency analysis measured as percentage change from baseline (400 ms pre-stimulus). B) Correction of multiple comparisons using cluster-based permutation for the time-frequency analysis showed a significant effect explained by a cluster that spans frequencies from around 10Hz to 30Hz in the duration from 0.5sec. to 1 sec. (n = 16, p = 0.0079).

**Supplementary Tables**

| **Sleep data** | **Mean ± SD** |
| --- | --- |
| TST (min) | 528.91 ± 37.38 |
| W (min) | 43.47 ± 39.05 |
| N1 (min) | 26.31 ± 14.20 |
| N2 (min) | 222.84 ± 39.85 |
| SWS (min) | 128.34 ± 44.23 |
| REM (min) | 95.09 ± 27.42 |

**Table S1. Sleep parameters.** Time spent in each sleep stages is reported in minutes and as percentage of total sleep time (mean ± SD). TST = total sleep time; W = wake; N1 = NREM Stage 1; N2 = NREM Stage 2; N3 = NREM Stage 3; REM = Rapid Eye Movement Sleep*.* N = 16.

| A) **LME model results** | | | | | |
| --- | --- | --- | --- | --- | --- |
| *Predictors* | *Model Estimates* | *CI* | *p* |  |  |
| *Intercept* | 3.03 | 2.68 – 3.38 | **<0.001** |  |  |
| *Cueing (Cued)* | -0.04 | -0.15 – 0.07 | 0.457 |  |  |
| *Baseline* | 0.43 | 0.37 – 0.49 | **<0.001** |  |  |
| *Session (2-Wk)* | -0.29 | -0.41 – -0.17 | **<0.001** |  |  |
| *Cueing (Cued) x Baseline* | -0.10 | -0.17 – -0.02 | **0.010** |  |  |
| *Session (2-Wk)* | 0.05 | -0.12 – 0.21 | 0.564 |  |  |
| B) **Post-hoc analysis** | | | | | |
|  | *Model Estimate* | *SE* | *df* | *t.stat* | *p* |
| *Baseline = -2* |  |  |  |  |  |
| *Uncued > Cued* | -0.18 | 0.09 | 1518 | 2.02 | **0.04** |
| *Baseline = -1* |  |  |  |  |  |
| *Uncued > Cued* | -0.08 | 0.06 | 1519 | 1.38 | **0.17** |
| *Baseline = 0* |  |  |  |  |  |
| *Uncued > Cued* | 0.02 | 0.04 | 1515 | -0.42 | 0.67 |
| *Baseline = 1* |  |  |  |  |  |
| *Uncued > Cued* | 0.11 | 0.06 | 1511 | -2.04 | **0.04** |
| *Baseline = 2* |  |  |  |  |  |
| *Uncued > Cued* | 0.21 | 0.09 | 1511 | -2.46 | **0.01** |

**Table S2.** A) Results of the mixed effect model examining the effect of Cueing (Cued/Uncued), Sessions (Session 48-H /Session 2-Wk) and pre-sleep Baseline on subjective arousal ratings. Model Estimates column lists regression coefficients. For each line, the reference value is indicated in brackets (e.g. Cueing (cued)). Notably the significant intercept indicates the predicted mean value of the model when all parameters and conditions are at reference level. B) post-hoc analysis of items with each mean baseline rating (-2 to 2), showing the basis of the interaction from (A).

|  | **Mean** | | **Standard Error (SE)** | |
| --- | --- | --- | --- | --- |
|  | **Cued** | **Uncued** | **Cued** | **Uncued** |
| **Session 1 (S1)** | 3.04 | 3.08 | 0.05 | 0.05 |
| **Session 2 (S2) – 48h after S1** | 2.97 | 3.04 | 0.06 | 0.05 |
| **Session 3 (S3) – 2 weeks after S1** | 2.83 | 2.85 | 0.06 | 0.05 |

**Table S3.** Descriptive statistic of the Arousal Rating Task.

| **Cued** | | | | | **Uncued** | | |
| --- | --- | --- | --- | --- | --- | --- | --- |
|  |  | Pearson’s  correlation | p-value | p-value  (FDR corr) | Pearson’s  correlation | p-value | p-value  (FDR corr) |
| Ratings S2 | Insula | 0.23 | 0.36 | 0.54 | 0.06 | 0.80 | 0.80 |
|  | OFC | 0.18 | 0.49 | 0.56 | 0.15 | 0.56 | 0.56 |
|  | sgACC | -0.01 | 0.97 | 0.97 | -0.20 | 0.44 | 0.66 |
|  | Amygdala | 0.10 | 0.70 | 0.70 | 0.38 | 0.12 | 0.36 |
| HRD | Insula | -0.09 | 0.75 | 0.75 | 0.33 | 0.25 | 0.33 |
|  | OFC | -0.25 | 0.38 | 0.38 | 0.36 | 0.20 | 0.27 |
|  | sgACC | -0.63 | 0.02 | 0.08 | 0.15 | 0.62 | 0.62 |
|  | Amygdala | 0.14 | 0.63 | 0.63 | 0.39 | 0.17 | 0.23 |

**Table S4. Correlations between mean parameter estimates, behavioural results, and**

**HRD.** Results of Pearson’s correlations between the mean parameter estimates for

each subject in our ROIs, behavioral results in S2, and heart rate deceleration

(HRD). Both the uncorrected and FDR-corrected p-values are reported.

| **Cued** | | | | | **Uncued** | | |
| --- | --- | --- | --- | --- | --- | --- | --- |
|  |  | Pearson’s  correlation | p-value | p-value  (FDR corr) | Pearson’s  correlation | p-value | p-value  (FDR corr) |
| HRD | TST (min) | 0.01 | 0.95 | 0.97 | -0.09 | 0.76 | 0.94 |
|  | N2 (%) | 0.21 | 0.44 | 0.97 | 0.06 | 0.85 | 0.94 |
|  | N3 (%) | -0.09 | 0.75 | 0.97 | -0.06 | 0.84 | 0.94 |
|  | REM (%) | 0.24 | 0.37 | 0.97 | 0.19 | 0.52 | 0.94 |
| HRD | Ratings S2 | 0.07 | 0.81 | 0.82 | 0.13 | 0.65 | 0.73 |

**Table S5. Correlations between HRD, sleep, and behavioural measurements**. Results

of Pearson’s correlations between HRD for cued and uncued image-sound

pairs and the percentage of time spent in each sleep stage, total sleep time, and

behavioural results in S2. Both the uncorrected and FDR-corrected p=values are

reported.

|  | AIC | BIC | Chisq | df | Pr(>Chisq) |
| --- | --- | --- | --- | --- | --- |
| Model 1 | 4260.83 | 4298.32 |  |  |  |
| Model 2 | 4001.07 | 4043.92 | 261.75 | 1 | 0.00 |
| Model 3 | 3996.48 | 4044.68 | 6.60 | 1 | 0.01 |

**Table S6. Model comparison.** AIC = Akaike Information Criterion; BIC = Bayesian Information Criterion are both measures of the relative quality of statistical models. Chisq = Chi-squared, test statistic for the LRT.

| **Slide No.** | **Description** | **Valence (M±SD)** | **Arousal (M±SD)** |
| --- | --- | --- | --- |
| 1112 | Snake | 4.71(1.70) | 4.60(2.44) |
| 1300 | PitBull | 3.55(1.78) | 6.79(1.84) |
| 1930 | Shark | 3.79(1.92) | 6.42(2.07) |
| 2095 | Toddler | 1.79(1.18) | 5.25(2.34) |
| 2717 | DrugAddict | 2.58(1.32) | 5.70(2.16) |
| 2799 | Funeral | 2.42(1.41) | 5.02(1.99) |
| 2800 | CryingBoy | 2.45(1.42) | 5.09(2.15) |
| 3016 | Mutilation | 1.90(1.31) | 5.82(2.44) |
| 3051 | Mutilation | 2.30(1.86) | 5.62(2.45) |
| 3068 | Mutilation | 1.80(1.56) | 6.77(2.49) |
| 3170 | BabyTumor | 1.46(1.01) | 7.21(1.99) |
| 3210 | Surgery | 4.49(1.91) | 5.39(1.91) |
| 3215 | BurnVictim | 2.51(1.32) | 5.44(2.16) |
| 3225 | Mutilation | 1.82(1.22) | 5.95(2.46) |
| 3230 | Dyingman | 2.02(1.30) | 5.41(2.21) |
| 3261 | Tumor | 1.82(1.34) | 5.75(2.64) |
| 3301 | InjuredChild | 1.80(1.28) | 5.21(2.26) |
| 6000 | Prison | 4.04(1.74) | 4.91(2.17) |
| 6021 | Assault | 2.21(1.51) | 6.06(2.38) |
| 6242 | Gang | 2.69(1.59) | 5.43(2.36) |
| 6315 | BeatenFem | 2.31(1.69) | 6.38(2.39) |
| 6550 | Attack | 2.73(2.38) | 7.09(1.98) |
| 6560 | Attack | 2.16(1.41) | 6.53(2.42) |
| 6834 | Police | 2.91(1.73) | 6.28(1.90) |
| 8485 | Fire | 2.73(1.62) | 6.46(2.10) |
| 9000 | Cemetery | 2.55(1.55) | 4.06(2.2.5) |
| 9041 | ScaredChild | 2.98(1.58) | 4.64(2.26) |
| 9050 | PlaneCrash | 2.42(1.61) | 6.36(1.97) |
| 9230 | OilFire | 3.89(1.58) | 5.77(2.36) |
| 9252 | DeadBody | 1.98(1.59) | 6.64(2.33) |
| 9253 | Mutilation | 2.00(1.19) | 5.53(2.40) |
| 9254 | Assault | 2.03(1.35) | 6.04(2.35) |
| 9301 | Toilet | 2.26(1.56) | 5.28(2.46) |
| 9320 | Vomit | 2.65(1.92) | 4.93(2.70) |
| 9405 | SlicedHead | 1.83(1.17) | 6.08(2.40) |
| 9410 | Soldier | 1.51(1.15) | 7.07(2.06) |
| 9420 | Soldier | 2.32(1.59) | 5.69(2.28) |
| 9423 | Assault | 2.61(1.51) | 5.66(2.15) |
| 9424 | Bomb | 2.87(1.62) | 5.78(2.12) |
| 9425 | Assault | 2.67(1.44) | 5.92(2.13) |
| 9429 | Assault | 2.68(1.26) | 5.63(2.04) |
| 9571 | Cat | 2.46(1.61) | 5.41(2.27) |
| 9582 | DentalExam | 4.18(2.28) | 5.29(2.21) |
| 9622 | Jet | 3.10(1.90) | 6.26(1.98) |
| 9630 | Bomb | 2.96(1.72) | 6.06(2.22) |
| 9800 | Skinhead | 2.04(1.57) | 6.05(2.71) |
| 9900 | CarAccident | 2.46(1.39) | 5.58(2.13) |
| 9921 | Fire | 2.04(1.47) | 6.52(1.94) |

**Table S7.** 48 standardized images taken from the IAPS database. Overall valence = Mean ± SD, 2.57 ± 0.75; overall arousal = Mean ± SD, 5.81 ± 0.68.
